# Telomere lengths in plants are correlated with flowering time variation

**DOI:** 10.1101/2020.07.04.188102

**Authors:** Jae Young Choi, Michael D. Purugganan

## Abstract

Telomeres are highly repetitive tandemly repeating DNA sequences found at chromosomal ends that protect chromosomes from deterioration during cell division. Using whole genome re-sequencing data, we found substantial natural intraspecific variation in telomere lengths in Arabidopsis thaliana, Oryza sativa (rice) and Zea mays (maize). Genome-wide association mapping in A. thaliana identifies a region that includes the telomerase reverse transcriptase (TERT) gene as underlying telomere length variation. TERT appears to exist in two haplotype groups (L and S), of which the L haplogroup allele shows evidence of a selective sweep in Arabidopsis. We find that telomere length is negatively correlated with flowering time variation not only in A. thaliana, but also in maize and rice, indicating a link between life history traits and chromosome integrity. We suggest that longer telomeres may be more adaptive in plants that have faster developmental rates (and therefore flower earlier), and that chromosomal structure itself is an adaptive trait associated with plant life history strategies.

## Introduction

Telomeres are a region of repetitive sequences that caps the end of eukaryotic chromosomes to protect them from deterioration (Greider and Blackburn, 1985). During DNA replication, failure to fill in terminal basepairs at the lagging strand leads to the “end-replication problem” (Olovnikov, 1973, 1971; Watson, 1972), resulting in the shortening of chromosome ends at each cell division and eventual loss of replicative capacity (Hayflick and Moorhead, 1961; van Deursen, 2014). To prevent this loss of chromosome termini, the ribonucleoprotein enzyme complex telomerase, whose core components consist of a telomerase reverse transcriptase (TERT) and RNA template (TER) (Osterhage and Friedman, 2009) binds to single-stranded telomeric DNA at the 3’ end and processively extends the telomere sequence (Wu et al., 2017). Other specialized telomere binding proteins are also recruited to prevent the telomere from being detected as damaged DNA (Fulcher et al., 2014).

Eukaryotic telomeres consist of a tandem repeat of TG-rich microsatellite sequences (Podlevsky and Chen, 2016). Between species, the core telomeric repeat sequence is conserved - for instance, vertebrates have the telomeric repeat TTAGGG (Meyne et al., 1989) while in most plants the sequence is TTTAGGG (Fajkus et al., 2005). The most noticeable telomere difference between organisms are in telomere lengths, which can be as short as 300 bps in yeast (Gatbonton et al., 2006) to 150 Kb in tobacco (Fajkus et al., 1995). Within species, telomere sequences also display substantial length variation, and several examples of telomere length polymorphisms and the underlying genes responsible for this variation have been identified in humans, yeast and C. elegans (Codd et al., 2013; Cook et al., 2016; Jones et al., 2012; Levy et al., 2010; Liti et al., 2009). In plants, variation in telomere lengths have also been observed between individuals (Burr et al., 1992; Fulcher et al., 2015; Maillet et al., 2006; Shakirov and Shippen, 2004), between organs (Kilian et al., 1995), and between cell types (González-García et al., 2015). Quantitative trait locus (QTL) studies in A. thaliana and maize have indicated that natural variation in telomere length is a heritable complex trait (Brown et al., 2011; Burr et al., 1992; Fulcher et al., 2015), although no specific genes have been identified.

A more puzzling question is what significance does natural variation in telomere lengths have for organisms? Telomere length variation could be neutral and result from random genetic drift or random stochasticity in the activity of the telomerase. Alternatively, telomere length differences could have fitness effects that are subject to natural selection, possibly due to their association with cellular senescence that has been implicated in controlling lifespan in yeast and animals (Aubert and Lansdorp, 2008; Kupiec, 2014). In mammals, for example, telomere shortening correlates with between-species differences in lifespans (Whittemore et al., 2019), suggesting telomeres are involved in the aging process (Aubert and Lansdorp, 2008). Indeed, it has been suggested that the aging trajectory of telomere lengths could be a product of optimization of a life-history tradeoff (Young, 2018). This is by no means universal, as in C. elegans no fitness differences or clear phenotypic consequences were associated with natural variation in telomere lengths (Cook et al., 2016).

While there is interest in the links between telomeres and life history traits (e.g., aging) in animals, comparatively little is known about how telomere length evolution impacts plant life history strategies. Aging in plants differs fundamentally from animals (Watson and Riha, 2011) and it is unclear whether the telomere-aging and evolution model are also applicable in plants. Indeed, no specific hypothesis have been put forth to explain natural telomere length variation in plants; whether telomeres have an effect on plant life history traits and are a target of natural selection remains an open question.

Here we describe the genetic basis and biological significance of natural telomere length variation in plants. Using whole genome sequence data, we determine the extent of telomere length variation in three plant species – Arabidopsis thaliana, Oryza sativa and Zea mays. We find that polymorphisms in the TERT gene is associated with natural telomere length variation in A. thaliana, and show that longer telomeres are found in plants that flower earlier. We propose a telomere-developmental rate model for plants wherein telomere length is an adaptive trait of individuals with specific life history strategies.

## Results

### Genome-wide variation in A. thaliana tandem repeats

Satellite DNA are repetitive sequences structured as arrays of DNA that are tandemly repeated in the genome, sometimes up to 106 copies. We examined genome-wide variation in satellite DNA repeat copy number in A. thaliana using the program k-Seek (Wei et al., 2018, 2014). k-Seek is an assembly-free method of identifying and quantifying k-mer repeats in unmapped short read sequence data, and k-mer counts are highly correlated with direct measurements of satellite repeat abundances (Wei et al., 2014).

We used whole genome re-sequencing data from the 1001 A. thaliana Genome Consortium project (Alonso-Blanco et al., 2016). We quantified genome-wide A. thaliana tandem repeat copy numbers by focusing on 483 individuals which were sequenced from leaves with identical protocols (designated as AraTha483; see Materials and Methods for details). The quantity of each k-mer sequence is presented as copies per 1× read depth after GC normalization (Flynn et al., 2017).

Adding up k-mer copy numbers, the median total length of tandem repeats per individuals is estimated at 341 Kb (Supplemental Figure 1). Across the population, individuals displayed over 25-fold differences in total tandem repeat lengths. The most abundant k-mer was the poly-A repeat, followed by the 7-mer AAACCCT (Fig. 1A). Some k-mers, such as the AC repeat, had a wide range of variation between individuals, with a range of 0 to 1,000s of copies. Our computationally based estimates were qualitatively concordant with direct estimates of repeat copy number that used Southern blot analysis to characterize 1- to 4-mer variation in a single A. thaliana ecotype (Depeiges et al., 1995).

**Figure 1.**
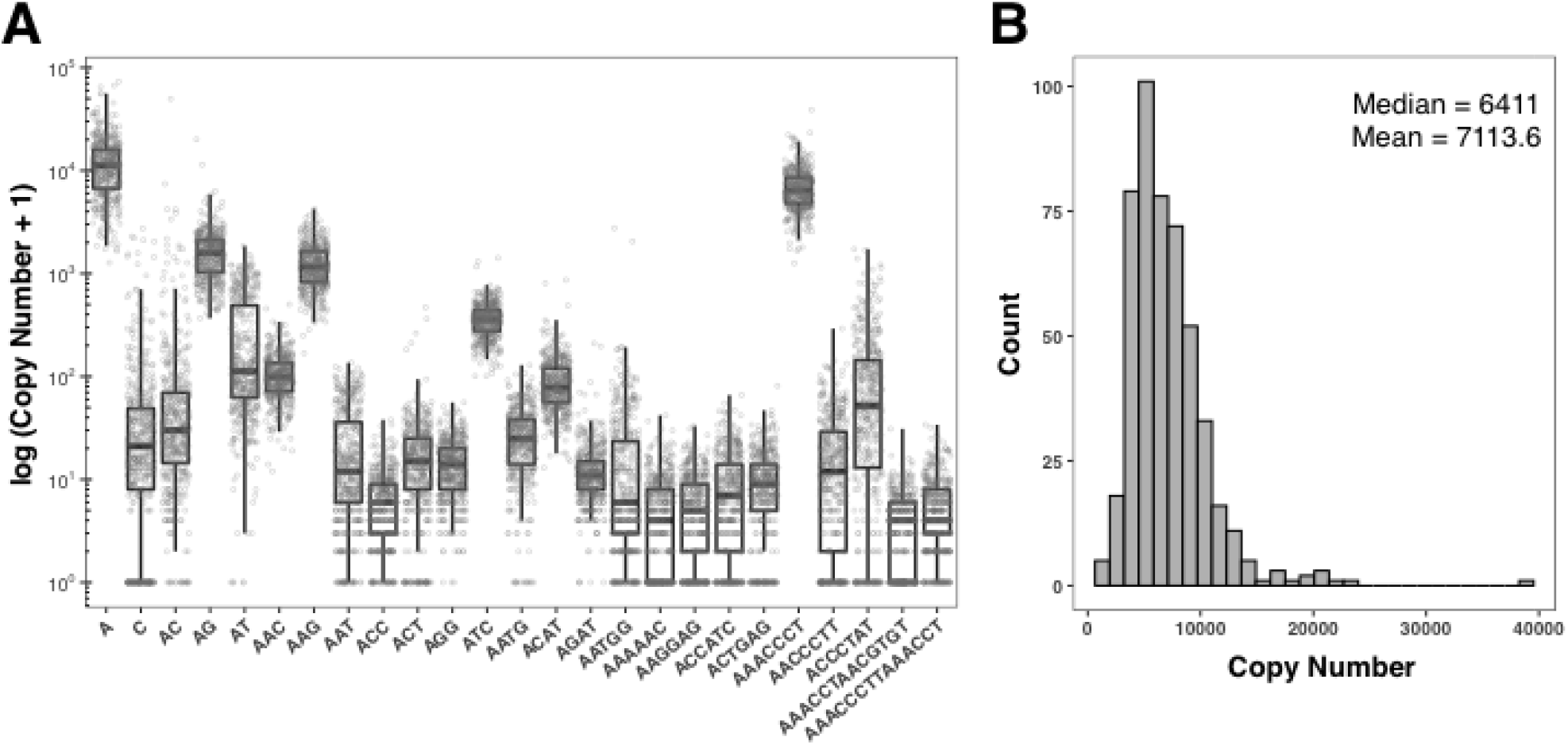
A. thaliana tandem repeat profile. Repeats were estimated from AraTha483 set. (A) Top 25 most abundant k-mers. K-mers are ordered alphabetically, then by size. (B) Distribution of estimates of telomere repeat copy number.

### A. thaliana telomere copy number variation and telomerase

The tandem repeat with the second highest abundance in the A. thaliana genome is the k-mer AAACCCT, which is the canonical telomere repeat sequence in plants [equivalent to the reported TTTAGGG telomere repeat] (Fajkus et al., 2005; Watson and Riha, 2010). There is a wide range in total copy numbers for the AAACCCT repeat, from 1,257 copies in ecotype Ler-1 to 38,850 copies in ecotype IP-Fel-2 (Supplemental Table 1), with a median of 6,411 and mean of 7,113.6 ± 161.1 copies (see Fig. 1B). We compared telomere repeat copy numbers inferred from k-Seek to a previous study that directly measured telomere lengths in various A. thaliana accessions using Southern blot analysis (Fulcher et al., 2015). Because the AraTha483 dataset had only 7 overlapping accessions with the Fulcher et al. dataset, we looked at data from a second set of 201 accessions (here on designated as AraTha201) that were sequenced with different protocols from the AraTha483 set. There were 53 accessions in common between AraTha201 and the Fulcher et al. samples, and we found a significant positive correlation in estimated telomere lengths from the two methods (Supplemental Fig 2; Pearson’s r = 0.61 and p = 1.26 × 10-6; Kendall’s tau = 0.189 and p = 0.046), suggesting that the k-mer approach is a valid approach to quantifying total telomere lengths.

We investigated whether natural variation in telomere length has a genetic basis, and using the AraTha483 set, we conducted genome wide association (GWAS) mapping of telomere copy number variation. We used the FarmCPU method for the GWAS analysis, which works well for identifying loci of complex traits that may be confounded with population structure (Liu et al., 2016). GWAS analysis revealed five genomic regions with single nucleotide polymorphisms (SNPs) significantly associated with telomere repeat copy number (Fig. 2A and see Supplemental Table 2 for SNP positions). The most significant SNP marker (p < 2.05 × 10-10) is located on chromosome 5 and found at the 3’ UTR of locus AT5G16850 (Fig. 2B) which is the telomerase reverse transcriptase (TERT) gene. The TERT gene is crucial in maintaining telomere lengths in A. thaliana (Fitzgerald et al., 1999) and other eukaryotes (Autexier and Lue, 2006). This SNP is also located in a quantitative trait locus (QTL) for telomere length previously identified in a recombinant inbred mapping study (Fulcher et al., 2015). The other four significant SNPs from the GWAS study were not in proximity to other known telomere regulating genes. However, two significant SNPs were also located in two QTL regions in chromosome 1 and chromosome 2 from the Fulcher et al. (2015) study (Fig. 2A).

**Figure 2.**
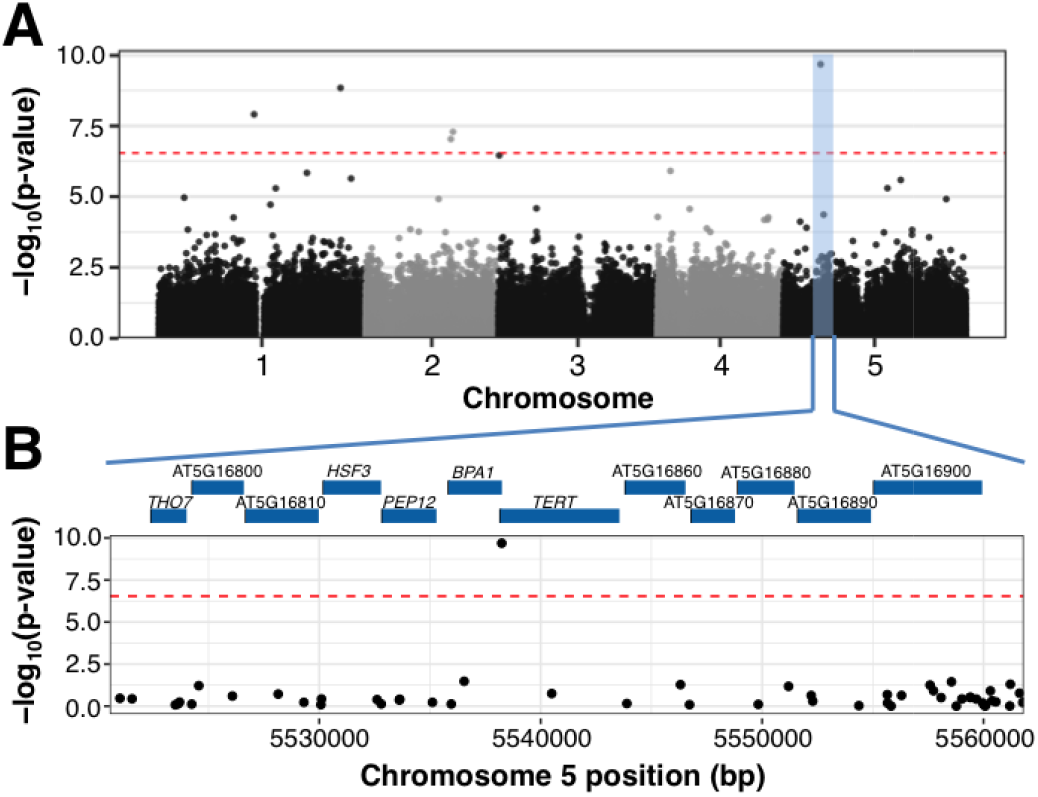
Genome wide association (GWAS) analysis of A. thaliana telomere length variation. Analysis was on the AraTha483 set. (A) Manhattan plot of the genome wide p-values testing association of telomere copy number using the FarmCPU approach. Red otted line indicates the Bonferroni-corrected significance threshold (α = 0.05). The region of the most significant of the five significant SNP regions is higlighted in blue. (B) Close-up of the region with the most significant GWAS SNP, with the genes in the region indicated.

### Population genetics of the A. thaliana TERT gene

Haplotype network reconstruction of the TERT gene in A. thaliana showed that TERT alleles are largely divided into two major haplotype groups (haplogroups) [see Fig. 3A]. Haplogroup L (Longer) differed from haplogroup S (Shorter) by 5 mutations at TERT (Fig. 3B), and individuals carrying haplogroup L had significantly higher telomere repeat copy numbers compared to haplogroup S (Mann Whitney U [MWU] test, p = 1.15 ×10-8; see Fig. 3C). The significantly higher telomere copy numbers for individuals with haplogroup L were also observed in the AraTha201 set (MWU test, p = 0.0098). In addition, telomere lengths of haplogroup L individuals that were directly estimated by Southern blot analysis (Fulcher et al. 2015) are also significantly longer than those from haplogroup S (MWU test, p = 0.009 and Fig. 3C).

**Figure 3.**
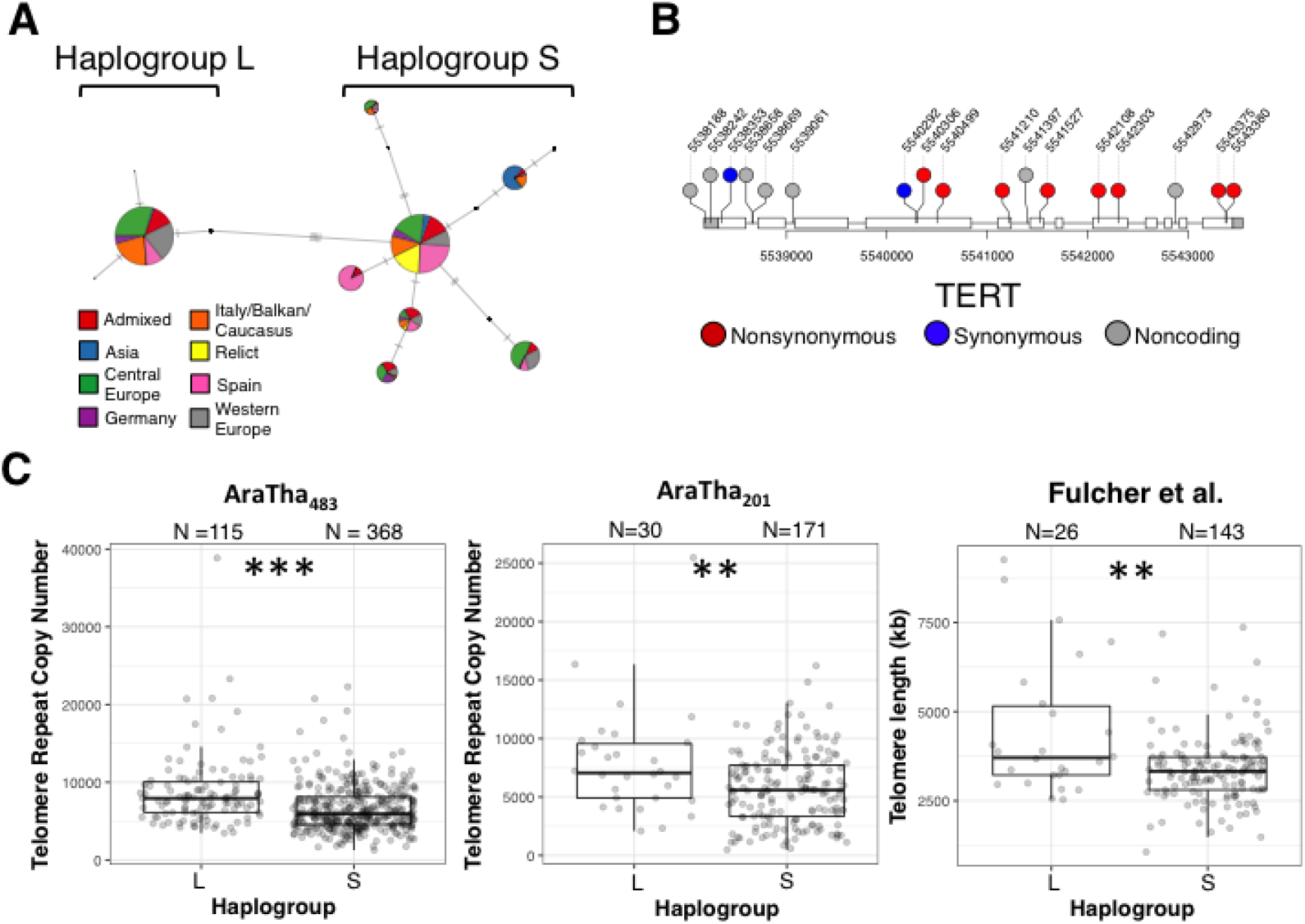
Haplotype analysis of the A. thaliana TERT gene. (A) Haplotype network of the TERT gene across the AraTha483 set. The haplogroup status that was designated from this study is indicated. Ancestry group status was taken from Alonso-Blanco et al. 2016. (B) TERT gene structure and the SNPs observed across the AraTha483 set. Haplogroup L specific mutations are indicated in the upper first row. (C) Comparison of telomere lengths between individuals carrying TERT haplogroup L or S. Comparisons were made from three different datasets. ** indicates p < 0.01 and *** indicates p < 0.001.

The TERT haplogroups were non-randomly distrib ted across geography. Haplogroup L was most common across individuals with Western European and Italy/Balkan/Caucasus ancestry, but was at low frequency in individuals with Asian and relic ancestry (Supplemental Table 3). Telomere copy number had significant negative correlations with both latitude and longitude (Fig. 4A), although a multiple linear regression model with both latitude and longitude showed only latitude as having a significant negative effect on telomere copy number (Supplemental Table 4). Overall, across the 1,135 samples from the 1001 A. thaliana Consortium samples, the frequency of haplogroup L was relatively low (15.1%), potentially explaining why previous attempts at GWAS mapping on telomere length variation failed to find significant associations (Fulcher et al., 2015). Further, differential frequency of TERT haplogroup with ancestry suggests that population structure could confound GWAS analysis (Atwell et al., 2010). Indeed using GWAS mixed linear models [i.e. MLM (Yu et al., 2006), CMLM (Zhang et al., 2010), MLMM (Segura et al., 2012), and SUPER (Wang et al., 2014)), the control for population structure effectively erased associations observed from the FarmCPU method (Supplemental Fig 3).

**Figure 4.**
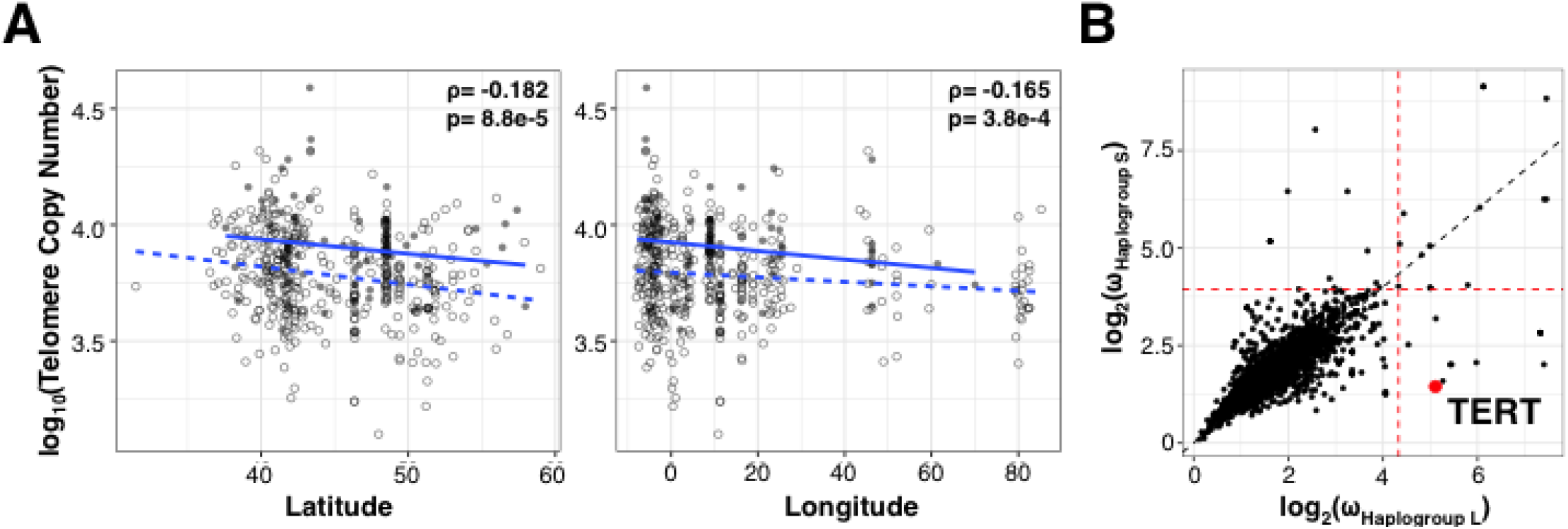
Evolution of the A. thaliana TERT gene. (A) Correlation between telomere copy number and latitude/longitude in the AraTha483 set. Closed and open circles represent individuals from haplogroup L and S, respectively. Dotted and solid lines represent line of best fit for haplogroup L and S samples, respectovely. The overall correlation (Spearman’s ρ) and p-value is indicated. (B) OmegaPlus-based selective sweep test statistics for individuals carrying TERT haplogroup L or S in the 1001 A. thaliana Genome Consortium samples. Red lines indicate the top 1% OmegaPlus statistic threshold.

The non-random distribution of TERT haplogroups with respect to geography and ancestry groups may be due to selection. To identify possible selective sweeps associated with TERT haplogroups, we marked the 1001 A. thaliana accessions based on which TERT haplogroup they carried, and applied OmegaPlus (Alachiotis et al., 2012; Kim and Nielsen, 2004) which uses linkage disequilibrium (LD) to detect selective sweeps in each group. We focused on chromosome 5 and looked for evidence of selection in 2,662 10-kb windows in L and S group accessions. Using a 1% empirical threshold we found evidence for selective sweeps at 38 genomic regions, one of which spanned the TERT gene, with selection only observed in individuals carrying TERT haplogroup L (Fig. 4B).

### A. thaliana telomere is associated with flowering time variation

The biased geographical distribution of telomere lengths and TERT genotypes suggest that length variation might have arisen as a geographical adaptation to specific environments. In A. thaliana, life history traits are often associated with geographic adaptation (Montesinos-Navarro et al., 2012; Stinchcombe et al., 2004), and we hypothesized that telomere length polymorphisms occurred as a response to adaptation to a specific life history strategy. We tested whether specific developmental traits associated with life history were correlated with variation in telomere length. We compared telomere copy number of the AraTha483 individuals to 7 different developmental traits and found significant negative correlations with 4 traits: day to flowering at 10°C (Spearman’s ◻ = −0.119, P < 0.009), day to flowering at 16°C (◻ = −0.173, p < 1.6 × 10-4), cauline leaf number (◻ = −0.125, p < 0.007), and rosette leaf number (◻ = −0.152, p < 0.001), and positive correlation with rosette branch number [◻ = 0.111, p < 0.03] (Fig. 5A). We also examined the direct telomere length measurements from Fulcher et al. (2015) and also found significant negative correlations with day to flowering at 10°C (◻ = −0.209, p < 0.004), day to flowering at 16°C (◻ = −0.213, p < 0.005), cauline leaf number (◻ = −0.216, p < 0.018), and rosette leaf number [◻ = −0.231, p < 0.014] (Supplemental Table 5).

**Figure 5.**
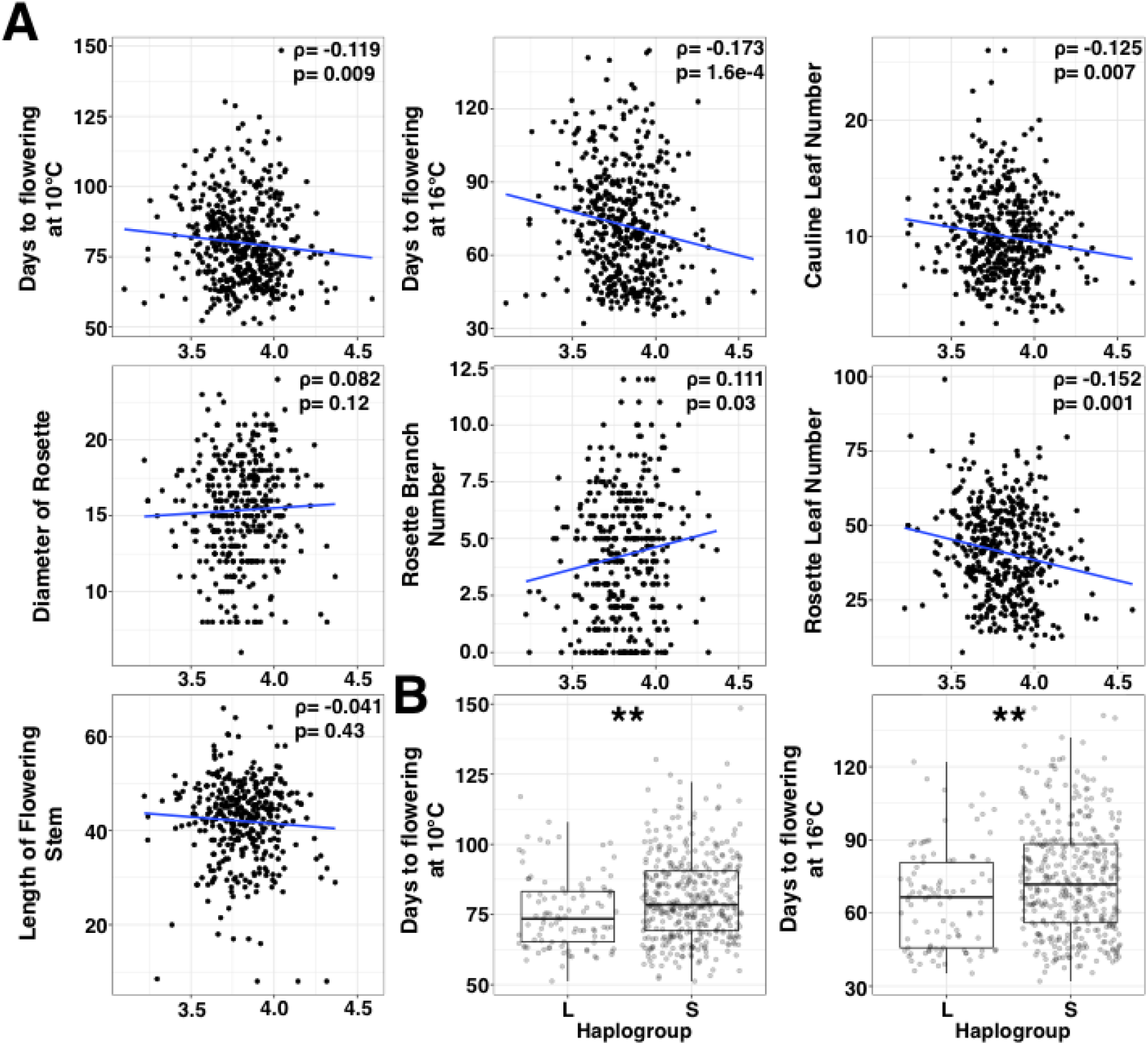
Association between telomere length and developmental traits in A. thaliana. (A) Correlation between telomere copy number and various A. thaliana life history traits for the AraTha483 set. Upper right corner shows the overall correlation (Spearman’s ρ) and significance. (B) Flowering time for individuals carrying TERT haplogroup L or S in the 1001 A. thaliana Genome Consortium samples. ** indicates p < 0.01.

To test whether these correlations were due simply to kinship/population structure, we fit a multiple linear regression model that included the first 4 axes of a principal component analysis of SNP variation. The results showed that telomere copy number was a significant negative predictor for the traits day to flowering at 16°C (p < 0.024), cauline leaf number (P < 0.034), and rosette leaf number (p < 0.003) even when accounting for population structure (Supplemental Table 6). It should be noted that in A. thaliana rosette leaf number is developmentally correlated with flowering time; together, these results suggest that telomere length is negatively associated with flowering time in this annual species, such that plants with longer telomeres flower earlier.

We expanded the analysis to the 1001 A. thaliana genome consortium samples by looking at the relationships between TERT haplogroup and developmental trait values (Fig. 5B). Results showed flowering time was the only trait that significantly differed between TERT haplogroups, with haplogroup L individuals flowering significantly earlier at both 10°C (MWU, p = 0.01) and 16°C (MWU, p = 0.0004).

Due to the significant associations of telomere length and TERT haplogroup with flowering time, we examined whether telomere regulating genes were in fact previously unrecognized flowering time QTLs (and vice versa). Using the AraTha483 individuals we conducted GWAS analysis on flowering time and compared the res lts to our GWAS analysis on telomere copy number (Fig. 2). Results showed there were no overlapping GWAS hits between the two traits, indicating that they had distinct genetic architectures (Supplemental Table 7 and Supplemental Figure 4A). Moreover, SNP markers in the TERT region were not significantly associated with flowering time (Supplemental Figure 4B). This suggests that while there is phenotypic correlation between telomere length and flowering time, this is not determined by common genes of pleiotropic effect.

### Flowering time is also negatively correlated with telomere copy number in rice and maize

The association between telomere length and flowering time was unexpected, but suggested individuals with different telomere lengths had contrasting life history strategies. We investigated if this correlation is found outside A. thaliana by examining the relationship between telomere length and flowering time in Oryza sativa and Zea mays. For each species we analyzed whole genome re-sequencing data from previous studies that also reported flowering time data (Flint-Garcia et al., 2005; Wang et al., 2018).

In rice (O. sativa) and maize (Z. mays) there was a wide variation in telomere copy number and, like A. thaliana, many of the differences appear to show population stratification (Supplemental Figure 5). We analyzed data for 2,952 rice varieties (Wang et al., 2018) and this species displayed the most significant differences between subpopulations, likely due to deep population structure in rice (Huang et al., 2012; Wang et al., 2018). Most rice varieties can be divided into japonica or indica subspecies (Wang et al., 2018), which possess significant genetic and physiological differentiation with each other (Zhao et al., 2011), and we analyzed each subpopulation separately (Fig. 6). In japonica, the temperate japonica (GJtmp) group had significantly higher telomere repeat copies than both subtropical (GJsubtrp) and tropical japonica (GJtrp) (MWU test, p = 0.0051 and 1.34 × 10-10 respectively). In indica rice, the subpopulation XI-1A (from East Asia) had significantly higher telomere copy numbers compared to subpopulation XI-1B (modern varieties of diverse origin), XI-2 (from South Asia), and XI-3 (from Southeast Asia) (MWU test, p = 0.0046, 2.38 × 10-20 and 1.83 × 10-18 respectively)[see Fig. 6].

**Figure 6.**
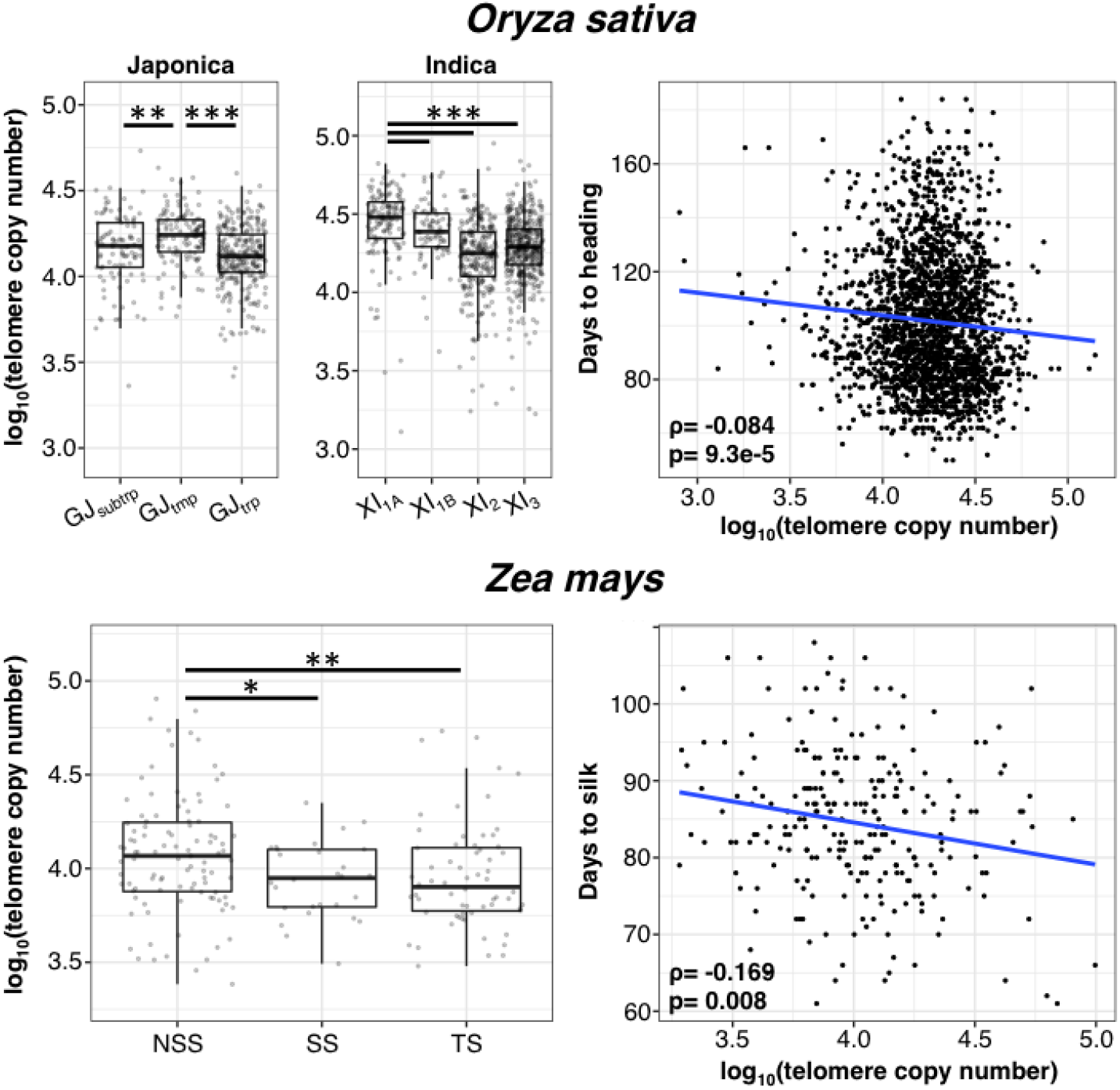
Association between telomere length and flowering time in rice (O. sativa) and maize (Z. mays). Telomere copy numbers per subpopulation are shown on left. * indicates p < 0.05, ** indicates p < 0.01, and *** indicates p < 0.001. Correlation between telomere copy number and flowering time is shown on right. For maize the days to silk measured from Aurora, NY at 2007 is shown but see Supplemental Table 8 for full results. Spearman ρ and p-values are shown in lower left of each plot.

In maize (Flint-Garcia et al., 2005) most varieties are genetically classified as either from non-stiff-stalk (NSS) and stiff-stalk (SS) populations from temperate regions (Liu et al., 2003), or from the tropical/subtropical (TS) population. Our analysis of 277 maize cultivars shows that NSS varieties had significantly higher telomere copy number than both SS and TS maize cultivars (MWU test, p = 0.0304 and 0.0065. respectively)[see Fig. 6]. Noticeably, in both rice and maize, the subpopulations with the highest telomere copy numbers were from temperate regions, and these had significantly higher abundance compared to varieties from subtropical or tropical regions.

Like in A. thaliana, we observe significant negative correlation between telomere copy number and flowering time in rice (◻ = −0.084, p < 9.3 × 10-5) [see Fig. 6]. In maize, the flowering time data was collected in 5 field locations in the United States over a 3 year period in some locations; there was data for a total of 9 fields/seasons (Zhao et al., 2006). In 6 cases, there was significant negative correlation of telomere repeat copy number and flowering time (◻ = −0.123 to −0.169, p < 0.008 to 0.045)[see Fig. 6 for one example], one was marginally non-significant (◻ = −0.130, p < 0.057) and one was still negative but non-significant (◻ = −0.057, p < 0.43) [Supplemental Table 8]. For both rice and maize a multiple linear model incorporating both telomere copy number and population structure as predictor variables still showed telomere length having a significantly negative effect on flowering time even after accounting for population stratification (p < 0.02 for rice and p < 0.033 for maize) [Supplemental Table 9].

GWAS analysis of telomere repeat copy number variation in rice and maize using FarmCPU showed significant SNP markers in the japonica rice, indica rice, and maize populations (Supplemental Figure 6). There were 16, 11, and 9 SNPs in indica rice, japonica rice, and maize, respectively, that were significant after Bonferroni correction (Supplemental Table 10). We identified 19 rice and maize orthologs of known telomere regulating genes (Supplemental Table 11) and compared their genomic positions to the GWAS significant SNP makers; none of the significant SNPs were in close proximity to these telomere regulating genes. We also examined the genetic architecture underlying flowering time variation in rice and maize, and like in A. thaliana we do not find any overlap in significant SNP positions for telomere length and flowering time variation (Supplemental Table 12).

## Discussion

Using whole genome re-sequencing data, we were able to computationally estimate total telomere length in individual plant genotypes by quantifying telomere repeat copy numbers. As expected, we find substantial intraspecific variation in genome-wid telomere lengths in plant species as diverse as A. thaliana, O. sativa and Z. mays. Interestingly, the genetic architecture of telomere length variation is distinct in these three species, and only in A. thaliana can we implicate a key telomere regulating gene – the telomerase TERT gene – in natural variation in telomere repeat copy number. The A. thaliana TERT gene is involved in telomere elongation (Shakirov and Shippen, 2004), and this locus also overlapped a large QTL region for telomere length identified from a previous recombinant inbred line mapping study (Fulcher et al., 2015). TERT has also been identified in a human GWAS mapping study showing an association with leukocyte telomere length variation (Codd et al., 2013).

The links between telomere lengths and organismal life history traits are tantalizing, especially since telomeres are linked to cellular senescence, aging and human disease. Despite its central role in chromosomal stability, the drivers of telomere length variation and their phenotypic consequences remain unclear. This is particularly relevant for plants, where telomere length variation is not easily connected to aging and senescence as observed in animals (Watson and Riha, 2011). In our analysis, we find that natural telomere length variation in three species is related to flowering time, one of the most crucial life history traits for plants. In Arabidopsis, rice and maize, individuals that had longer telomeres flowered earlier, and finding this correlation in three distinct species suggest that this relationship may be widespread. Indeed, we also observed this negative correlation between flowering time and telomere length in the polyploid Brassica napus, but this relationship was not significant in this species (unpublished results).

The correlation between telomere length and flowering time could arise from pleiotropic effects either of telomere regulating or flowering time genes. Our GWAS results, however, do not show any overlap in significant peaks between these two traits, indicating that they are controlled by distinct loci, and suggests that there is unlikely to be a direct causal genetic connection between telomere length and flowering time. What may drive this correlation are telomere length polymorphisms occurring as a response to adaptation to specific plant life history strategies. The link between telomere length and flowering time variation may reside in differences in plant developmental rates. In maize, for example, faster rates of cell differentiation in the shoot apical meristem is observed with earlier flowering times (Bilinski et al., 2018; Leiboff et al., 2015), and telomerase is most active in differentiating tissues such as the meristem (Fitzgerald et al., 1996; Riha et al., 1998). We theorize that in life history strategies associated with early flowering, individuals with longer telomeres have a selective advantage due to greater stability of chromosomal ends as meristematic cells go through more rapid division and differentiation (Huffman et al., 2000; Kazda et al., 2012).

Indeed, there is evidence for a selective sweep at the A. thaliana TERT haplogroup L (which is associated with longer telomeres), supporting the hypothesis that the negative correlation between telomere length and flowering time is driven by adaptive evolution. Adaptation may also explain the significant latitudinal cline of telomere length variation in Arabidopsis, and we also find longer telomeres associated with other aspects of the spring cycling life history strategy of this ruderal species such as germination in response to cold (Supplementary Fig. 7). Moreover, longer telomere lengths are found in temperate-adapted varieties of rice (temperate japonica) and maize (Non-Stiff Stalk and Stiff Stalk maize), which also flower significantly earlier in their growing seasons compared to tropical/subtropical varieties (Supplementary Fig. 8).

There has been interest in identifying effects of genome size and structure on life history traits such as flowering time (Meagher and Vassiliadis, 2005). In maize, for example, genome size is positively correlated with flowering time (Jian et al., 2017) and changes in repetitive DNA sequences are associated with altitudinal adaptation (Bilinski et al., 2018). The negative correlation between genome size, repetitive DNA content and cellular growth rate has been advanced as a plausible explanation for this phenomenon (Bilinski et al., 2018; Tenaillon et al., 2016). These studies as well as our results on telomere variation suggest that variation in life history strategy can indirectly influence chromosome and genome structure via selection. This opens up future areas of inquiry, including determining how widespread is this phenomenon, the relationship of telomere length and cell differentiation rate in plants, details of any selective advantage of telomere length in different life histories, and the precise molecular genetic mechanisms underlying telomere length polymorphisms in plant species.

## Materials and Methods

### Analyzed genome sequences

We obtained whole genome resequencing data for A. thaliana, B. napus, O. sativa and Z. mays from previous published studies. For each species we analyzed sequence batches that would minimize the technical differences between individuals. This involved analyzing genome sequences form the same tissue type, sequenced on the same sequencing platform, the same sequencing library preparation, and the same sequencing read length. Under this guideline we analyzed the following genome sequences from each species:

Arabidopsis thaliana: Genome sequences were obtained from the 1001 A. thaliana genome consortium (Alonso-Blanco et al. 2016) and available at the NCBI SRA under the identifier SRP056687. Each of the 1,135 samples was genome sequenced using a different tissue for DNA extraction, differing sequencing read length, and under various sequencing platforms. Because of this we grouped samples that had the same genome sequencing origins and analyzed the three most highly represented groups. This included the first group (designated as AraTha483) with 483 individuals that were prepared using leaf tissue, genome sequenced as 2×100bp read length and using the Illumina HiSeq 2000 platform. The second group (designated as AraTha201) consisting of 201 individuals that were prepared using leaf tissue, genome sequenced as 2×101bp read length and using the Illumina HiSeq 2000 platform.
Oryza sativa: Genome sequencing data from (Wang et al. 2018) were obtained at the NCBI SRA under the identifier PRJEB6180. Since all sequenced samples were prepared from similar developmental time points (i.e. leaf tissue) and genome sequenced as 2×83bp using the Illumina HiSeq 2000 platform, we categorized all 3,000 samples as a single group and analyzed them together. Samples with greater than 5× genome coverage were only used.
Zea mays: We analyzed the “282” panel of (Flint-Garcia et al. 2005), which aim to select varities to capture the genetic diversity of maize. The 282 panel has been sequenced twice over the years but to minimize the potential differences arising from sequencing platforms we analyzed the most recent sequencing batch that had resequenced the 282 panel to a higher depth using 2×150bp and Illumina HiSeq Ten X platform (Bukowski et al. 2018). The data was obtained from the NCBI SRA under the identifier PRJNA389800.

### Identification and quantifying tandem repeats

Initially the sequencing reads were subjected to quality control using the BBTools suite (https://jgi.doe.gov/data-and-tools/bbtools/). We used the bbduk.sh script ver. 37.66 from BBTools using the parameters minlen=25 qtrim=rl trimq=10 ktrim=r k=25 mink=11 hdist=1 tpe tbo to trim off sequencing adapters and low quality sequences from the reads.

The quality controlled unmapped reads were used to quantify the tandem repeat landscape of an individuals’ genome using the k-Seek method (Wei et al. 2018, 2014). Briefly this method identifies the short tandem repeat sequences (k-mers) in the raw sequencing reads using a hash table approach. k-Seek breaks up a read into smaller fragments to build a hash table consisting of the fragmented sequences and its frequencies across the read. The hash table is then used to identify the shortest k-mer motif that is tandemly repeated in a given sequencing read. k-Seek can identify k-mers of length 1 to 20 bps and the k-mer must be a tandem repeat covering at least 50 bps of a read. The method allows a single nucleotide mismatch for a given repeating k-mer. After identifying and quantifying the repeating k-mer, the offset (i.e. AAC, ACA, and CAA are all considered the same repeat) and reverse complement (i.e. AAC and GTT are considered the same repeat) of the k-mers are complied for summing up the k-mer copies across the entire sequencing library. k-Seek in the end, identifies the total copy number for a tandemly repeating k-mer in a sample of an unmapped genome sequencing library.

While k-Seek has been shown to be highly accurate in identifying tandem repeats from the short read genome sequences (Wei et al. 2014), the PCR based library preparation that is usually used for generating the sequencing is known to have a bias in underrepresenting high and low GC regions of the genome (Benjamini and Speed 2012). To account for this bias we implemented the method of (Flynn et al. 2017) to correct for differences in GC content affecting the copy number count between k-mers repeats with differing GC content. Reads are mapped against the reference genome to first calculate the mean insert size using the program bamPEFragmentSize from the deeptools ver. 3.3.0 package (Ramírez et al. 2016). The insert size was used for calculating the GC content of a given position in the genome, which was defined as the proportion of G or C bases of a given position plus the downstream fragment length of the library (Benjamini and Speed 2012). The alignment was then used to calculate the average coverage of each GC content. We used bwa-mem version 0.7.16a-r1181 (Li 2013) with default parameters to align the paired end reads to the reference genome. The average coverage per GC content is then used to calculate a correction factor of (Benjamini and Speed 2012) and applied to the k-mer counts. We used the scripts from (Flynn et al. 2017) (https://github.com/jmf422/Daphnia-MA-lines/tree/master/GC_correction) that implements the entire process.

For the A. thaliana resequencing data we used the reference genome TAIR10 from The Arabidopsis Information Resource (TAIR) and implemented the method of (Flynn et al. 2017) to correct for GC content bias and genome coverage between A. thaliana genome sequencing samples. The genome sequencing for O. sativa, and Z. mays, however, were not ideal for implementing the GC content correction method. For O. sativa, the samples were sequenced across multiple runs suggesting any differences in the sequencing run should also be implemented in the correction. While for Z. mays the genome coverage was relatively low (on average ~5×) indicating a coverage based method of correction would not be ideal. Hence, for these three species we only analyzed the telomere repeat and for each sample its telomere count was divided by the average genome wide coverage to account for differences in sequencing coverage between samples. The per sample average coverage was obtained from Supplementary Data 2 of (Wang et al. 2018) for O. sativa and for Z. mays it was calculated using bedtools ver. 2.25.0 (Quinlan and Hall 2010) genomecov program.

### Genome wide association study

SNP variant files were obtained from the original studies that generated the genome resequencing data and conducted the SNP calls. Specifically, for A. thaliana the population VCF file was downloaded from the 1001 genomes project website (https://1001genomes.org/), for O. sativa the population VCF was downloaded from the 3000 rice genome projects’ snp-seek website (https://snp-seek.irri.org/) (Mansueto et al. 2017), and for Z. mays the population VCF was downloaded from Gigascience Database (http://dx.doi.org/10.5524/100339) which is associated with (Bukowski et al. 2018).

The SNP files were initially filtered to exclude polymorphic sites that had more then 10% of the individuals with a missing genotype and filtered out sites with less then 5% minor allele frequency. We then conducted a linkage disequilibrium (LD) based pruning to remove polymorphic sites. The VCF files were converted to PLINK format using vcftools version 0.1.15 (Danecek et al. 2011) and the program plink ver. 1.9 (Chang et al. 2015) was used with the parameter --indep-pairwise 100 5 0.5, which scans the file in 100 variant count windows while shifting the window in 5 variants and pruning pairs of variants that have a r2 greater then 0.5.

The LD pruned PLINK file was converted to a HAPMAP format to be used for the GWAS analysis using the program GAPIT (Tang et al. 2016). We took the log10 of the telomere copy number to transform the distribution. For detecting SNPs significantly associating with a phenotype we used the FarmCPU algorithm (Liu et al. 2016), which is a mixed linear model (MLM) incorporating population structure and kinship but is robust to false positive and negative associations then other MLM GWA algorithms. We used four principle components to model the underlying population structure.

Orthologs of A. thaliana telomere regulating genes were found in the rice and maize gene annotation using the program Orthofinder ver 2.3.12 (Emms and Kelly 2019, 2015).

### A. thaliana TERT gene analysis

The SNPs for the A. thaliana TERT region were extracted using vcftools and missing genotypes were imputed and phased using Beagle version 5.0 (Browning and Browning 2016). The haplotype network of the TERT region was then reconstructed using the R (R Core Team 2016) pegas (Paradis 2010) and VcfR (Knaus and Grünwald 2017) package. The hamming distance between haplotypes was used for constructing a minimum spanning tree. Effects of each SNP were determined through the program snpeff (Cingolani et al. 2012).

Evidence of selective sweep were examined using the OmegaPlus method (Alachiotis et al. 2012). We extracted SNPs from chromosome 5 to exclude polymorphic sites that had more then 10% of the individuals with a missing genotype and filtered out sites with less then 5% minor allele frequency. This filtered SNP file was used for imputing missing genotypes using Beagle. The data was divided into individuals belonging to haplogroup L or S and resulting VCF file was used for the OmegaPlus ver. 3.0.3 program. We executed the program with -grid 2697 so that each grid would correspond to roughly 10,000 bp, and additional parameters -minwin 5000 -maxwin 3000000 -no-singletons.

### Plant phenotype analysis

The phenotypes that were used for associating with telomere lengths were obtained from previous studies. For A. thaliana the various developmental traits were obtained from Arapheno (https://arapheno.1001genomes.org/) (Seren et al. 2017) with the phenotype names FT10 (days to flowering at 10°C), FT16 (days to flowering at 16°C), CL (cauline axillary branch number), RL (leaf number), Length (stem length), RBN (primary branch number), and Diameter (flower diameter). Seed germination response to vernalization was obtained from (Martínez-Berdeja et al. 2020), specifically from the 2nd principal component of Fig. 1 of that study.

For rice we obtained phenotype data that were measured as part of the 3000 rice genome project (Sanciangco et al. 2018; Wang et al. 2018). Dataset includes 32 rice traits but we only analyzed the flowering time data, which was measured by estimating the number of days at which 80% of the plants were fully headed (code HDG_80HEAD). The data is available from https://doi.org/10.7910/DVN/HGRSJG.

For maize we obtained phenotype data from the Buckler-Goodman association panel, which consists of 57 different traits measured across 16 different environments. Phenotype file (traitMatrix_maize282NAM_v15-130212.txt) was downloaded from Panzea (https://www.panzea.org/phenotypes) and we only analyzed the days to silk trait (code GDDDaystoSilk).

Association between the telomere length and plant phenotypes were conducted in R. The multiple linear regression analysis was conducted using the lm function and population structure information was obtained from the four principle components that was used in the GWAS analysis.

## Acknowledgement

We thank Kevin Wei with assistance in using the k-Seek method and discussions on satellite sequence evolution, and Jullien Flynn and Ian Caldas for assistance in implementing the GC-bias correction method. We also thank members of the Purugganan laboratory, especially Simon (Niels) Groen for helpful discussions. This work was supported in part by grants from the National Science Foundation Plant Genome Research Program (IOS-1546218) and the Zegar Family Foundation (A16-0051) to M.D.P.

