## Supplementary figures and images for "Telomere lengths in plants are correlated with flowering time variation"

### Supplemental Fig. 1

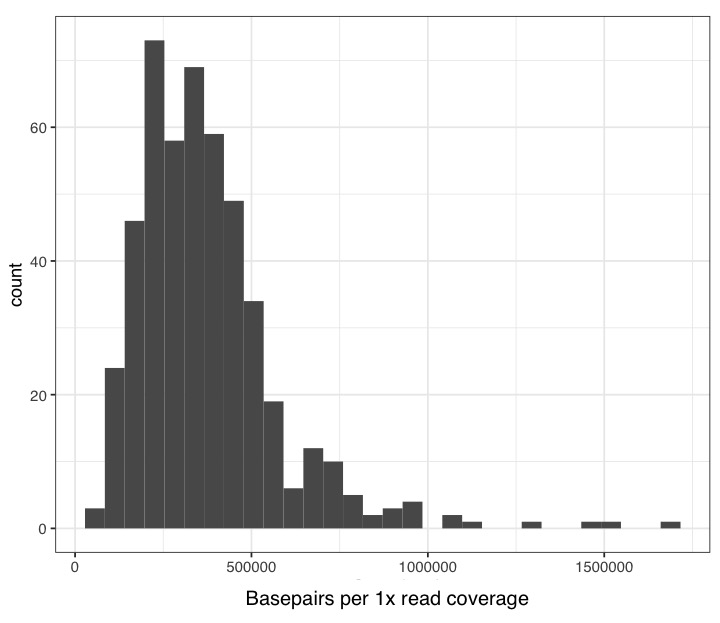

### Supplemental Fig. 2

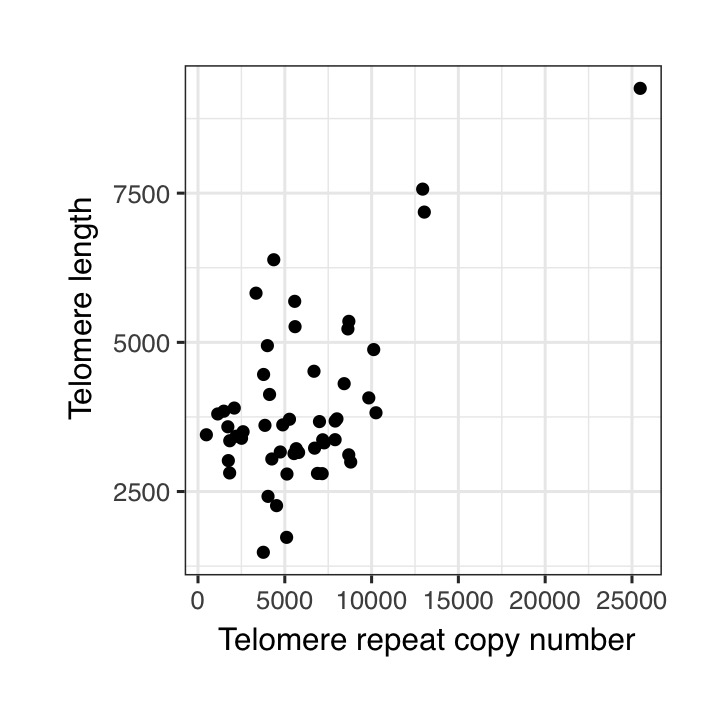

### Supplemental Fig. 3

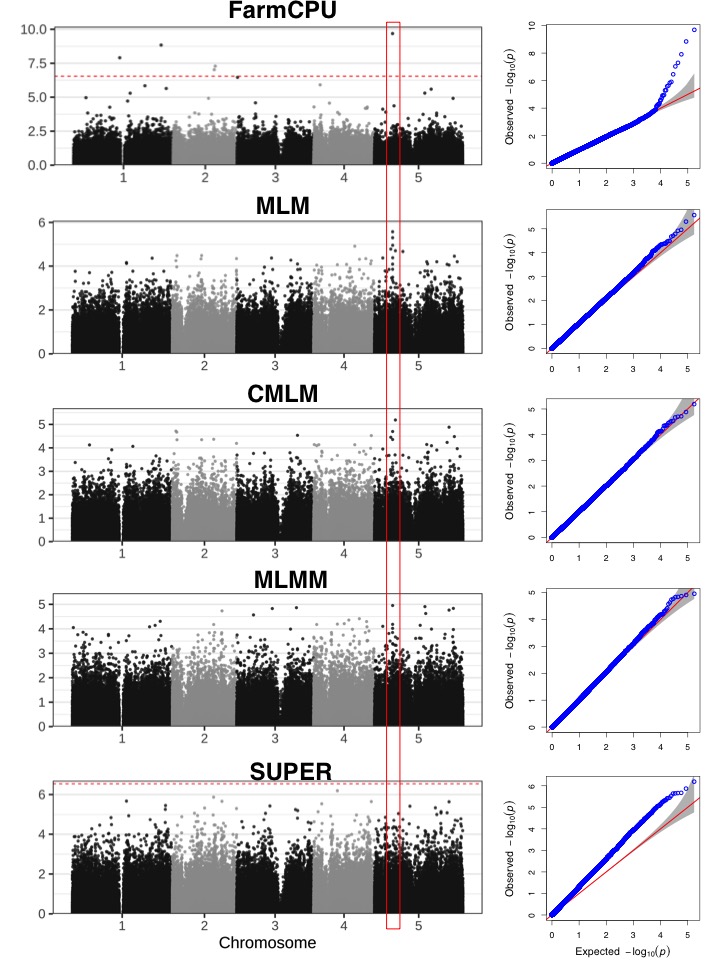

### Supplemental Fig. 4

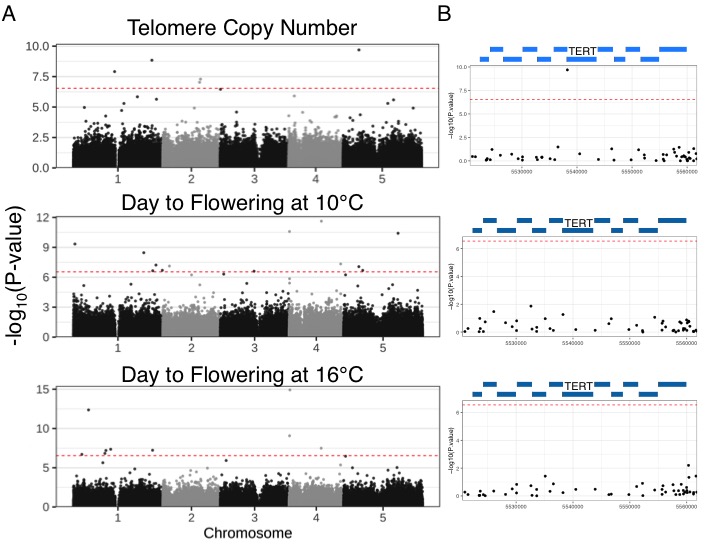

### Supplemental Fig. 5

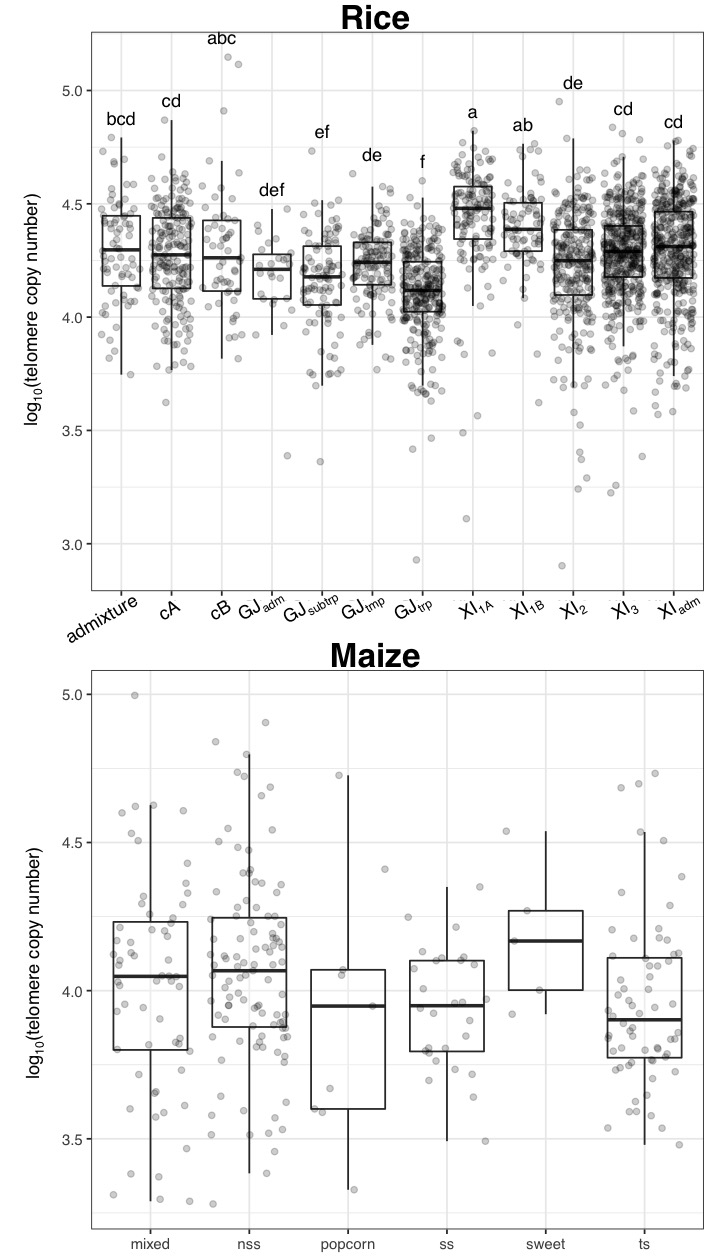

### Supplemental Fig. 6

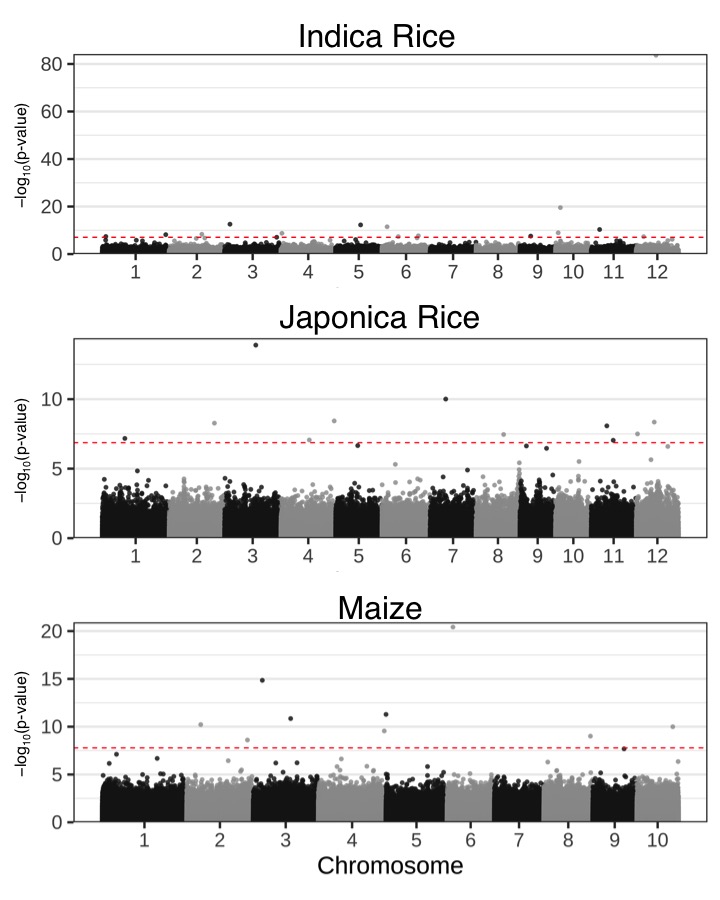

### Supplemental Fig. 7

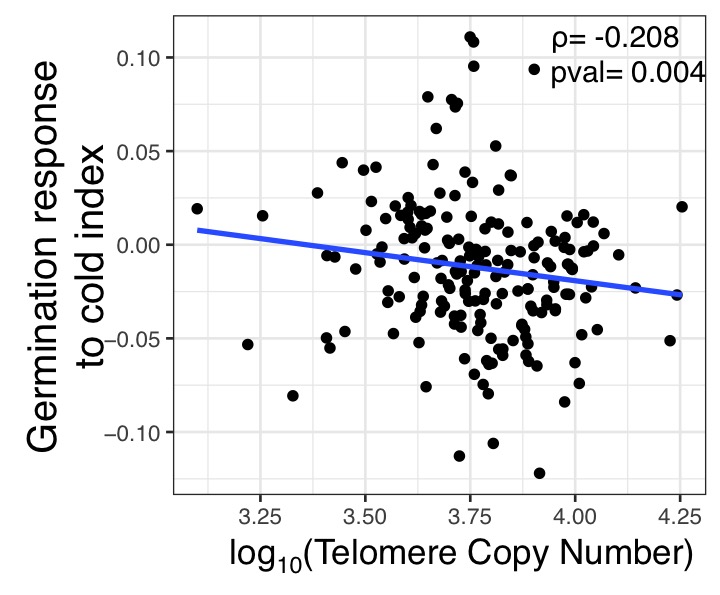

### Supplemental Fig. 8

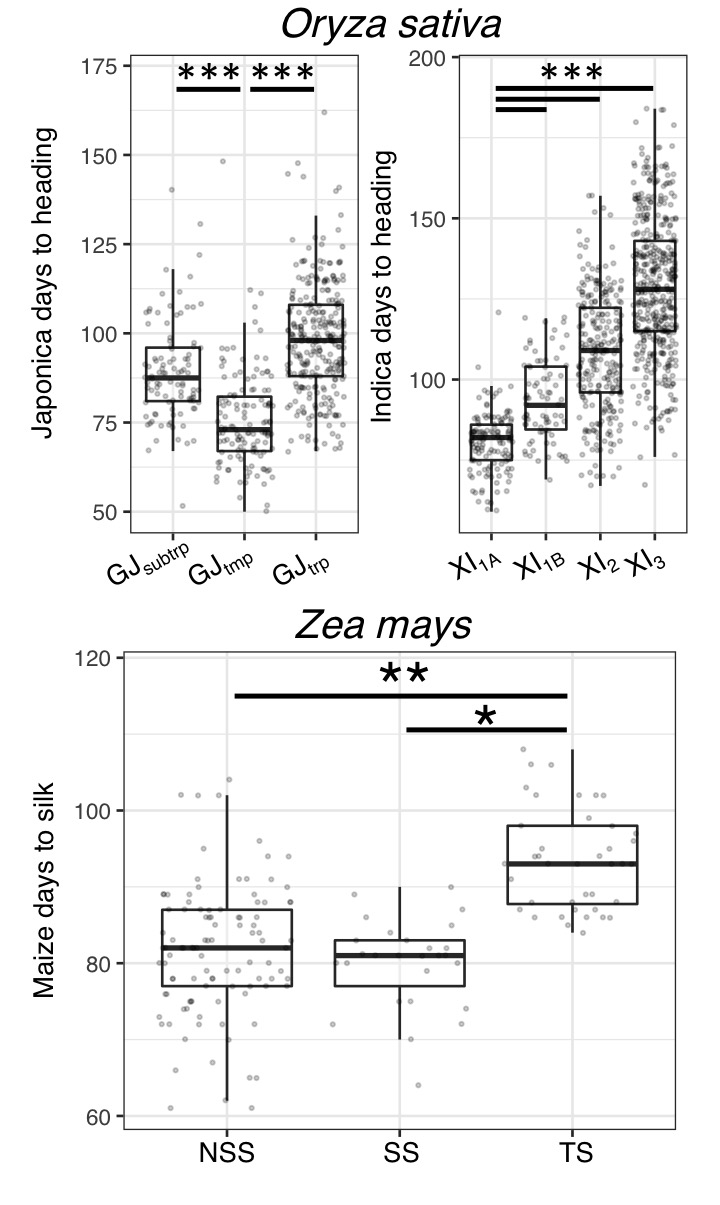
